# A rationalized definition of general tumor suppressor microRNAs excludes miR-34a

**DOI:** 10.1101/2021.02.11.430795

**Authors:** Sophie Mockly, Élisabeth Houbron, Hervé Seitz

## Abstract

While several microRNAs (miRNAs) have been proposed to act as tumor suppressors, a consensual definition of tumor suppressing miRNAs is still missing. Similarly to coding genes, we propose that tumor suppressor miRNAs must show evidence of genetic or epigenetic inactivation in cancers, and exhibit an anti-tumorigenic *(e.g.,* anti-proliferative) activity under endogenous expression levels. Here we observe that this definition excludes the most extensively studied tumor suppressor candidate miRNA, miR-34a. In analyzable cancer types, miR-34a does not appear to be down-regulated in primary tumors relatively to normal adjacent tissues. Deletion of *miR-34a* is occasionally found in human cancers, but it does not seem to be driven by an anti-tumorigenic activity of the miRNA, since it is not observed upon smaller, *miR-34a-specific* alterations. Its anti-proliferative action was observed upon large, supra-physiological transfection of synthetic miR-34a in cultured cells, and our data indicates that endogenous miR-34a levels do not have such an effect. Our results therefore argue against a general tumor suppressive function for miR-34a, providing an explanation to the lack of efficiency of synthetic miR-34a administration against solid tumors.

## Introduction

Tumor suppressors are genes whose activity antagonizes tumorigenesis. Consequently, they are frequently silenced, either by germline-inherited or somatic mutation, or otherwise inactivated, in cancers [1]. Mechanistically, tumor suppressors typically mediate cellular environment-induced inhibition of cell proliferation, therefore exhibiting anti-proliferative activity under their natural expression levels: a gene displaying cytotoxic or cytostatic activity only when inappropriately overexpressed is therefore excluded from that definition [2].

miRNAs are small regulatory RNAs, guiding their effector proteins to specific target RNAs, which are repressed by various mechanisms (target RNA degradation and translational inhibition) [3]. Targets are recognized by sequence complementarity, with most targets bearing a perfect match to the miRNA “seed” (nt 2–7) [4]. Such a short binding motif makes miRNA/target binding poorly specific, and more than 60% of human genes are predicted to be targeted by at least one miRNA [5]. Because such gene regulators can act in signal transduction cascades, they may participate in tumor-suppressive pathways. A consensual definition for “tumor suppressor miRNAs” is still lacking, with some tentative definitions being based on miRNA down-regulation in cancer cells [6], on the targets’ annotation [7], or both [8]. We rather propose to follow the initial definition of tumor suppressors [2], considering that there is no reason to particularize miRNAs among other types of tumor suppressors. We thus advocate for the following definition of tumor suppressor miRNAs: **(i)** there is evidence for their frequent inactivation in cancer (either by genetic or epigenetic alteration; potentially only in specific cancer types); and **(ii)** they inhibit tumorigenesis *(e.g.,* by repressing cell proliferation) under their endogenous expression level, rather than upon unrealistic overexpression.

We applied this definition to interrogate the status of the most highly-studied tumor suppressor candidate miRNA, miR-34a. It is a member of the miR-34 family, comprising six members in human and in mouse: miR-34a, miR-34b, miR-34c, miR-449a, miR-449b and miR-449c (Supplementary Figure S1). The three miR-34a/b/c subfamily members are transcriptionally controlled by the p53 tumor suppressor, which suggested that these miRNAs could participate in the tumor suppressive activity of the p53 network [9, 10, 11, 12, 13, 14, 15]. Indeed, the miR-34a member is down-regulated or lost in various cancer models (tumor samples or transformed cell lines) relatively to normal samples [9, 11, 16, 10, 14, 17, 18, 19, 20]. This observation suggested that the inactivation of *miR-34a* is involved in tumorigenesis, and that other family members *(miR-34b* and *c, miR-449a, b* and *c*) could not compensate for this loss. miR-34a is therefore widely perceived as a general tumor suppressor, whose inactivation is involved in a variety of cancer types [21]. Yet *miR-34a*^−/−^, *miR-34b*^−/−^, *miR-34c*^−/−^triple knock-out mice do not exhibit obvious defects in p53-dependent proliferation control or in tumor suppression [22]. And, while pre-clinical studies in mice gave encouraging results (reviewed in [23, 24]), administration of a synthetic miR-34a to human patients with solid tumors failed to repress tumor growth reproducibly [25]. An alternate administration regimen (allowing increased drug exposure) did not clearly improve clinical outcomes, while triggering poorly-understood, severe adverse effects [24].

## Materials and Methods

### Analysis of *miR-34a* expression and integrity in human cancers

miRNA expression data was downloaded from the GDC portal on April 29, 2021. Cancer types where at least 10 cases were available (with Small RNA-Seq data from normal solid tissue and primary tumor for each case) were selected, and depth-normalized read counts were compared between normal tissue and tumor for each case. The heatmap shown on Figure 1**A** shows the median log-ratio between tumor and normal tissue, with non-significant changes (calculated with the Wilcoxon test, FDR-adjusted for multiple hypothesis testing) being colored in white.

**Figure 1.**
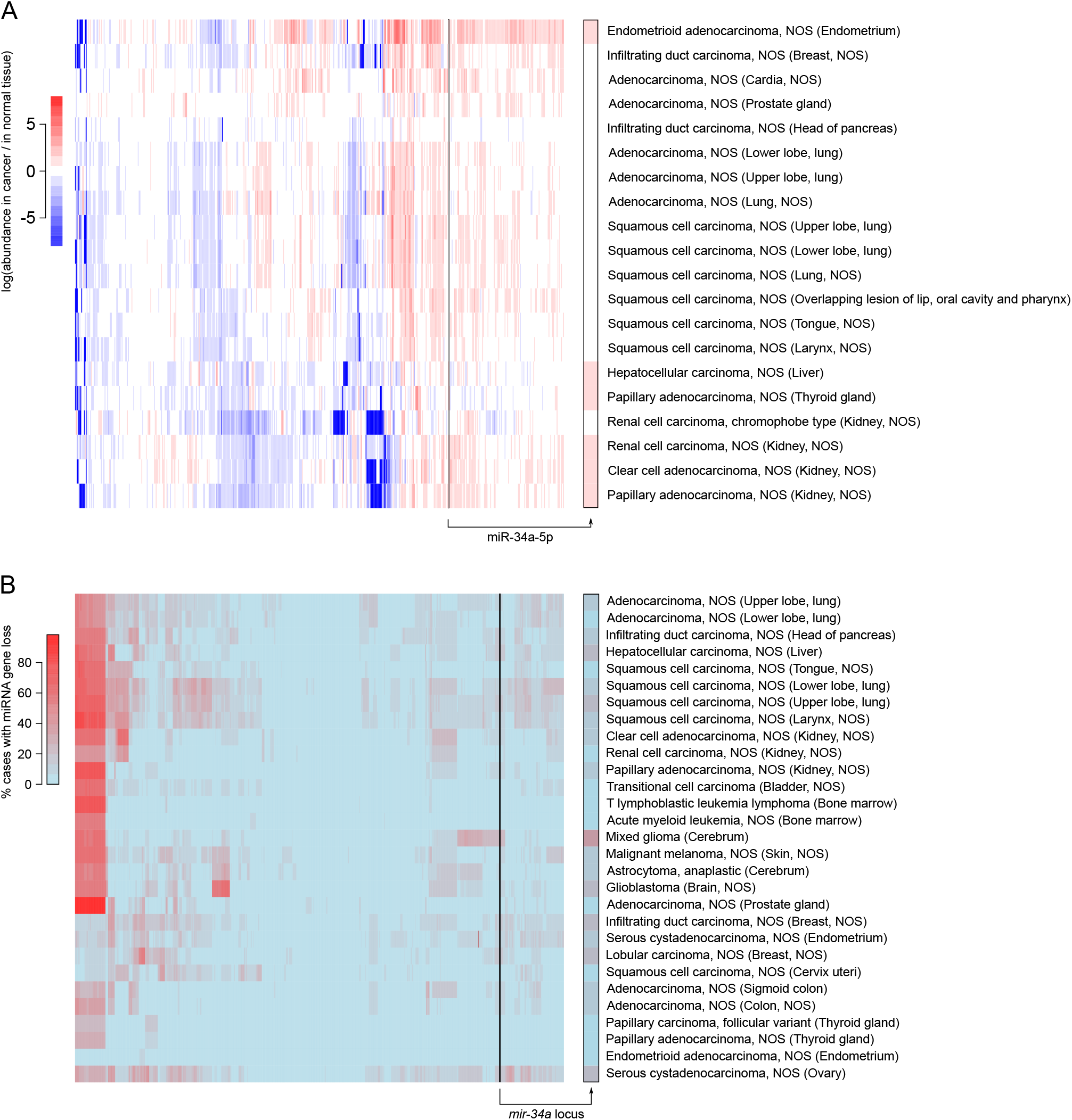
*mir-34a* is not generally down-regulated or lost in cancers. **A.** miRNA abundance (normalized by the number of mapped miRNA reads) was compared between primary tumors and normal adjacent tissues. Only cancer types for which at least 10 cases were analyzed have been considered (n=20 cancer types; rows), and miRNAs with a null variance across cancer types were excluded (remaining: n=545 miRNAs; columns). For each miRNA/cancer type pair, the heatmap shows its median fold-change across all cases, with non-significant changes (FDR ≥ 0.05) being shown in white. log(fold-changes) larger than +8 or smaller than −8 were set to +8 or −8 respectively, for graphical clarity. **B.** Only cancer types for which at least 100 cases were analyzed have been considered (n=29 cancer types; rows), and miRNA genes whose ploidy could not be assessed were excluded (remaining: n=1,686 miRNAs; columns). For each miRNA/cancer type pair, the heatmap shows the percentage of cases with monoallelic or biallelic loss of the miRNA gene. **Both panels:** the column showing miR-34a data is magnified on the right margin (framed in black). “NOS”: not otherwise specified.

miRNA gene ploidy data was downloaded from the GDC portal on March 4, 2021. Erroneous miRNA gene coordinates were corrected using information from miRBase. For the heatmap shown on Figure 1**B**, the percentage of cases with miRNA gene loss (either homo- or heterozygous) was evaluated for each miRNA, selecting cancer types where ploidy was determined in at least 100 cases. miRNA sequence variation data was downloaded from the GDC portal on February 24, 2021. SNP location was intersected with miRNA hairpin and mature miRNA coordinates from miRBase (as well as with miRNA seed coordinates, defined as nt 2–7 of the mature miRNA). For the heatmaps shown on Supplementary Figure S2, the percentage of cases with sequence variations in miRNA genes (hairpin, mature or seed sequences) is displayed, selecting cancer types with at least 100 analyzed cases.

For each of these heatmaps, miRNAs and cancer types were clustered with the heatmap.2 command with the **R** software.

### CRISPR/Cas9-mediated mutagenesis

Four sgRNAs were designed using CRISPOR (http://crispor.tefor.net/ [26]) to target each side of the human pre-mir-34a sequence, and cloned into an expression plasmid for *S. pyogenes* Cas9 (pSpCas9(BB)-2A-GFP plasmid (PX458), a gift from Feng Zhang [27]; Addgene plasmid #48138; http://n2t.net/addgene:48138; RRID:Addgene_48138). Targeting efficiency of each plasmid was estimated by Sanger sequencing of the targeted locus in transfected HCT-116 cells, and analyzed with the Synthego ICE Analysis online tool (https://ice.synthego.com/#/). Mutagenesis was performed using the most efficient sgRNA sequence on each side of the targeted locus (AAGCTCTTCT-GCGCCACGGT**GGG** and GCCGGTCCACGGCATCCGGA**GGG**; PAM sequences in bold; also see Supplementary Figure S5).

HCT-116 (ATCC^®^ cat. #CCL247) and HAP1 (Horizon Discovery cat. #C631) cells were grown till 80% confluency and transfected with the two plasmids (15 μg each) following the protocol for Lipofectamine 2000 Transfection Reagent (Thermo Fisher Scientific). After 24 hours, Cas9-GFP-expressing single cells were isolated in 96-well plates by flow cytometry on a BD FACSMelody (Becton Dickinson), then grown for 10 days. Homozygous wild-type and mutant clones were first tested by discriminative PCRs (with primer pairs ACTTCTAGGGCAGTATACTTGCT and GCT-GTGAGTGTTTCTTTGGC; and TCCTCCCCACATTTCCTTCT and GCAAACTTCTCCCAGC-CAAA), and eventually validated by Sanger sequencing of their *miR-34a* locus. For the HAP1 cell line, mutagenesis efficiency was so high that we were unable to isolate wild-type clones after cotransfection of sgRNA-carrying PX458 plasmids. Wild-type clones were therefore generated by transfection of HAP1 cells with a plasmid expressing SpCas9-HF1 variant but no sgRNA (the VP12 plasmid, a gift from Keith Joung [28]; Addgene plasmid #72247; http://n2t.net/addgene:72247; RRID:Addgene_72247), and went through the same isolation and selection process as mutant clones.

### RNA extraction

Cells plated in 10 cm Petri dishes were lysed and scrapped in 6 mL ice-cold TRIzol™ Reagent (Invitrogen) added directly to the culture dish after removal of the growth medium, and mixed with 1.2 mL of water-saturated chloroform. Samples were homogenized by vigorous shaking for 1 min and centrifuged for 5 min at 12,000 g and 4°C to allow phase separation. The aqueous phase was transfered in a new tube and mixed with 3 mL isopropanol for precipitation. After a 10 min incubation at room temperature, samples were centrifuged for 10 min at 12,000 g and 4°C and the supernatant was removed. The RNA pellet was washed with 6 mL of 70% ethanol and samples were centrifuged for 5 min at 12,000 g and 4°C. After complete removal of ethanol, the RNA pellet was resuspended in 20 μL RNase-free water and the quantity of total RNA was determined by spectrophotometry on a NanoDrop ND-1000.

### Small RNA-Seq

Total RNA of each cell line was extracted 48 h after seeding and quality was assessed on electrophoretic spectra from a Fragment Analyzer (Agilent), analyzed with the PROSize software (v. 3.0.1.6). Libraries were prepared using NEXTflex™ Small RNA-Seq Kit v3 (Bioo Scientific) following the manufacturer’s instructions. Libraries were verified by DNA quantification using Fragment Analyzer (kit High Sensitivity NGS), and by qPCR (ROCHE Light Cycler 480). Libraries were sequenced on Illumina NovaSeq 6000 using NovaSeq Reagent Kit (100 cycles). RNA quality assessment, library preparation, validation and sequencing were performed by the MGX sequencing facility.

Adapters ended with 4 randomized nucleotides in order to reduce ligation biases. Because of the sequencing design, the adapter sequence (5’ GTTCAGAGTTCTACAGTCCGACGATCNNNN 3’) appears at the beginning of the read sequence, and the final 4 nucleotides of the read are the initial randomized nucleotides of the other adapter, whose other nucleotides are not read. Hence small RNA reads can be extracted from the fastq files with the following command:

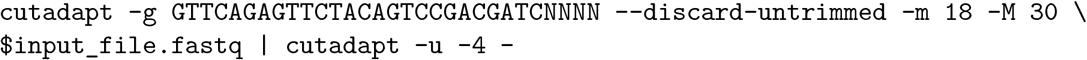

### Cell transfection

Cells were transfected 24 hours after seeding either with a control duplex, siRNA against eGFP: 5’-GGCAAGCUGACCCUGAAGUdTdT-3’ / 5’-ACUUCAGGGUCAGCUUGCCdTdT-3’ or with a hsa-miR-34a mimic duplex: 5’-P-UGGCAGUGUCUUAGCUGGUUGUU-3’ / 5’-P-CAAUCAGCAAGUAUACUGCCCUA-3’ according to the protocol for Lipofectamine 2000 Transfection Reagent (Thermo Fisher Scientific).

### Proliferation assays

Because the mere procedure of isolating and selecting mutated clones may artifactually select clones with exceptionally high proliferation rates, we applied the same isolation and selection procedure to wild-type clones, and we measured proliferation rates on several independent wild-type and mutant clones. Each cell line was seeded in 96-well plates (Figure 3**C**: in 4 replicates at 3 × 10^3^ cells/well per time point; Figures 4**A** and **B**: in 6 replicates at 6 × 10^3^ cells/well). From 24 hours after cell seeding or transfection, to 3 days later, the number of living cells was determined twice a day by CellTiter-Glo Luminescent Cell Viability Assay (Promega) according to the manufacturer’s protocol and recorded with a TriStar LB 941 (Berthold Technologies). Linear regression of log-transformed cell counts relative to time and genotype (in **R** syntax: `log-transformed cell counts ~ time * genotype`) or transfected duplex identity (`log-transformed cell counts ~ time * duplex identity`) was used to measure doubling time and to estimate the significance of the effect of genotype or transfected duplex.

For Figure 3**D** and **E**, doxorubicin (Sigma-Aldrich) was diluted in molecular biology-grade water and 5-fluorouracil (5-FU) (Sigma-Aldrich) diluted in dimethyl sulfoxide (SigmaAldrich). In a preliminary experiment, half-maximal inhibitory concentration (IC50) was estimated after 72 h drug exposure: 7 × 10^-8^ M and 8 × 10^-6^ M for doxorubicin and 5-FU respectively. Cell lines were seeded in 3 replicates per drug concentration at 2.5 × 10^3^ cells/well in 96-well plates. After 24 hours, culture medium was replaced with drug-containing medium (concentration range centered on the IC50 with 2.5× increments), or solvant-containing medium for untreated controls, and the number of living cells was determined 72 h later by CellTiterGlo Luminescent Cell Viability Assay (Promega). Cell counts were normalized to the mean cell number in untreated controls. Normalized cell number was fitted to an asymptotic model for each clone to assess the significance of the effect of genotype (using an analysis of variance to compare a model not informed by clone genotype, to a genotype-informed model).

### Measurement of apoptosis

HCT-116 cells were seeded in 6-well plates in 3 replicates at 9 × 10^4^ cells/well per condition. 72 hours after cell transfection, the number of apoptotic, dead and live cells was determined by FITC Annexin V/Dead Cell Apoptosis Kit with FITC annexin V and PI for flow cytometry (Invitrogen cat. #V13242), according to the manufacturer’s protocol. Cells were analyzed by flow cytometry on a MACSQuant analyzer (Miltenyi) using the blue laser excitation (488 nm) with a 525/50BP filter. 10,000 singlet cells were measured per replicate, and apoptotic, dead and live populations were defined by FITC and PI thresholds pre-established with non-stained and mono-stained controls, and counted with the FlowJo Software (BD Biosciences).

### miRNA quantification by RT-ddPCR

Reverse transcription of a specific miRNA in HCT-116 cells was performed on 10 ng total RNA using the TaqMan microRNA Reverse Transcription Kit (Thermo Fisher Scientific) in a total volume of 15 μL, according to the manufacturer’s protocol, with miRNA-specific RT primers from the TaqMan MicroRNA Assay Kit (assay IDs for hsa-miR-34a-5p and miR-21b-5p are respectively 000426 and 000397). In order to ensure a precise ddPCR quantification, with similar numbers of positive and negative droplets in each sample, cDNA dilution factor was optimized for each experimental condition (cDNAs for miR-21 quantification were diluted 10 ×; cDNAs for miR-34 quantification were diluted: 100× for 1 nM-transfected samples, 1000 × for 10 nM-transfected samples, undiluted for 0 nM-transfected samples and for samples shown in Figure 4**D**). ddPCR amplification of the cDNA was performed on 1.33 μL of each cDNA dilution combined with 1 μL of miRNA-specific 20X TaqMan MicroRNA Reagent containing probes and primers for amplification from the TaqMan MicroRNA Assay Kit (Thermo Fisher Scientific), 10 μL of 2X ddPCR Supermix for probes (no dUTP) (Bio-Rad), and 7.67 μL of molecular biology-grade water. Droplets were generated, thermal cycled and detected by the QX200 Droplet Digital PCR System (Bio-Rad) according to the ddPCR Supermix protocol and manufacturer’s instructions. Data were extracted using QuantaSoft Pro Software (Bio-Rad).

### Statistical analyses of RT-ddPCR data

miR-34a and miR-21 quantification was performed in 3 independent replicates, and cDNA counts were converted into numbers of miRNA molecules per ng RNA, considering dilution factors at each step in the RT-ddPCR process. Significance of the effect of transfected dose (for Figure 4**C**), doxorubicin treatment (for Figure 4**D**) and time (for both panels) was assessed by two-way ANOVA without interaction. Post-hoc pairwise t-tests were performed whenever the ANOVA found a significant effect for an explanatory variable.

### Data and script availability

Deep-sequencing data has been deposited at SRA and linked to BioProject number PRJNA695193. Scripts, raw, intermediate and final data files are available at https://github.com/HKeyHKey/Mockly_et_al_2022 and at https://www.igh.cnrs.fr/en/research/departments/genetics-development/systemic-impact-of-small-regulatory-rnas#programmes-informatiques/. In particular, flow cytometry raw data has been deposited at https://github.com/HKeyHKey/Mockly_et_al_2022/tree/main/Suppl_Figure_8/Flow_cytometry_raw_data.

## Results

### No evidence for *miR-34a* loss or inactivation in cancers

It is now possible to compare miRNA levels between tumors and normal adjacent tissues on a large collection of human cases which passed stringent, standardized quality controls [29], allowing a rigorous assessment of miR-34a expression in tumorigenesis. Selecting every cancer type where miRNA expression is available for primary tumor and normal adjacent tissue, in at least 10 studied cases (n=20 cancer types), we did not find any cancer type where miR-34a was significantly down-regulated (Figure 1**A**). Hence in this collection of cancer types, human primary tumors do not tend to under-express miR-34a, contradicting the notion that genetic or epigenetic silencing of *miR-34a* could participate in tumorigenesis.

Accordingly, genetic alterations affecting *miR-34a* are very rare in cancer: focusing on every cancer type for which gene-level copy number was measured in at least 100 cases (n=29 cancer types), we did not observe any tendency for the loss of *miR-34a* relatively to other miRNA genes (see Figure 1**B**). Similarly, we did not find any evidence for the selective mutation of the pre-miR-34a hairpin precursor sequence, mature miR-34a or the miR-34a seed in cancers (n=30 analyzed cancer types; Supplementary Figure S2). In contrast to *miR-34a,* 105 miRNA loci tend to be frequently lost in 19 cancer types (red area at the top left corner of the heatmap in Figure 1**B**; listed in Supplementary Table S1): these miRNAs are more convincing tumor suppressor candidates than *miR-34a* in this respect.

It could be argued that *miR-34a* inactivation by itself is insufficient to contribute to tumorigenesis, while it may play a role in a sensitized context, where additional, cooperative mutations may reveal the oncogenicity of miR-34a down-regulation. In that case, *miR-34a* inactivation could be enriched in just a subset of highly mutated cancers, and it would not be visible in the analyses shown in Figure 1 and Supplementary Figure S2. Yet, stratifying cases by cancer grade, we did not observe any tendency for the most aggressive tumors to inactivate *miR-34a* (Supplementary Figure S3), indicating that even the most sensitized tumors do not show any evidence of *miR-34a* inactivation.

Similarly, it is conceivable that miR-34a plays a tumor suppressive role only in the presence of functional p53, and the frequent mutation of p53 in the samples analyzed in Figure 1 may have obscured its behaviour in p53^+/+^ tumors. But the selective analysis of cancer cases without any mutation in *p53* gives a very similar result, without miR-34a being down-regulated in any analyzed cancer type (see Figure 2).

**Figure 2:**
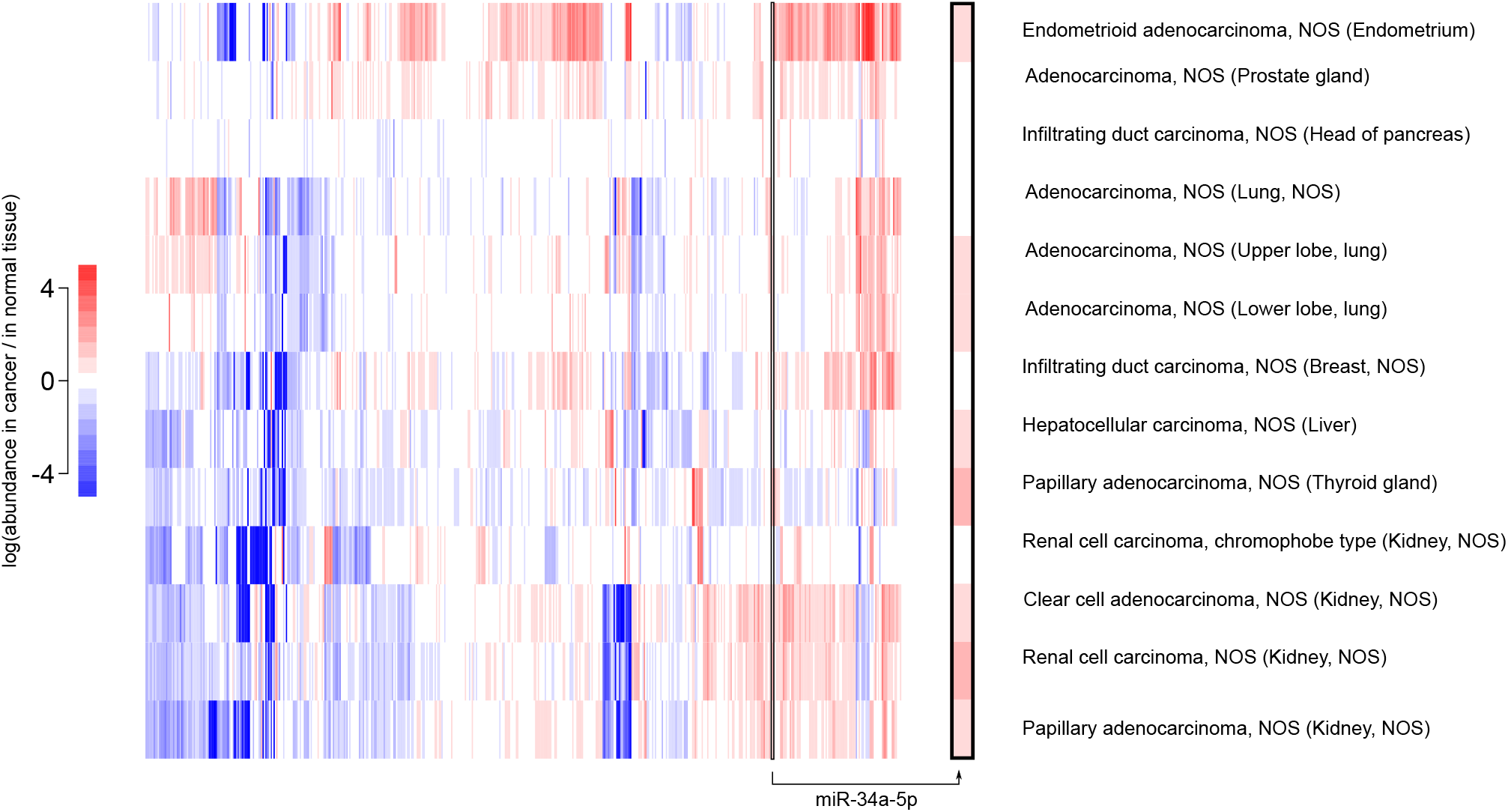
No evidence for *mir-34a* inactivation in tumors with an intact *p53* gene. Cancer samples analyzed in Figure 1**A** were stratified by the mutation status of the *p53* gene, and only cases without any detected mutation in *p53* were selected here (also selecting cancer types with at least 10 cases after this selection). Same conventions than in Figure 1**A**. miRNA abundance (normalized by the number of mapped miRNA reads) was compared between primary tumors and normal adjacent tissues. The column showing miR-34a data is magnified on the right margin (framed in black). log(fold-changes) larger than +5 or smaller than −5 were set to +5 or −5 respectively, for graphical clarity. “NOS”: not otherwise specified.

Hence the loss or mutation of *miR-34a* does not appear to be enriched in cancer. We note that *miR-34a* is located on cytogenetic band 1p36, which is often altered in a wide variety of cancers. But our analyses suggest that the inactivation of *miR-34a* is not the actual driver for deletion selection – and because a convincing tumor suppressor is already known at 1p36 (the *CHD5* gene [30]), we propose that the occasional deletion of *miR-34a* in cancer is rather a consequence of its genomic proximity with such a real tumor suppressor. Accordingly, whenever a limited region of consistent deletion could be mapped in 1p36, that region excludes *miR-34a* (with the only exception of myelodysplastic syndromes, but with low experimental support): see Supplementary Figure S4.

### The reported anti-proliferative action of miR-34a is artifactual

miR-34a has also been considered a tumor suppressor candidate on the basis of the apparent antiproliferative activity of miR-34 family miRNAs. Numerous studies in cultured cell lines indeed showed that miR-34 transfection inhibits cell proliferation, either by slowing down cell division or by increasing cell death [16, 9, 11, 12, 13, 14, 15]. But miRNA over-expression generates false positives, raising the possibility that this reported anti-proliferative role is artifactual [31]. We thus deleted the *miR-34a* gene in HCT-116 cells, where it has been proposed to be anti-proliferative by several independent studies [9, 11, 14] (mutagenesis strategy in Supplementary Figure S5). Deletion of the *miR-34a* locus eliminated 94% of the expression of the whole miR-34 family (Figure 3**A** and **B**). Our results do not show any significant difference in the growth rate of *miR-34a^−/−^* and wild-type clones (Figure 3**C**). We also prepared *miR-34a^−^* clones from the human haploid HAP1 cell line, where miR-34a is also not anti-proliferative (it is even slightly pro-proliferative; Supplementary Figure S6). It could be argued that *miR-34a* does not inhibit cell proliferation in unstressed conditions, while being anti-proliferative upon genotoxic stress. But we also failed to observe significant differences between wild-type and mutant clones under doxorubicin or 5-fluoro-uracil treatment (Figure 3**D** and **E**).

**Figure 3:**
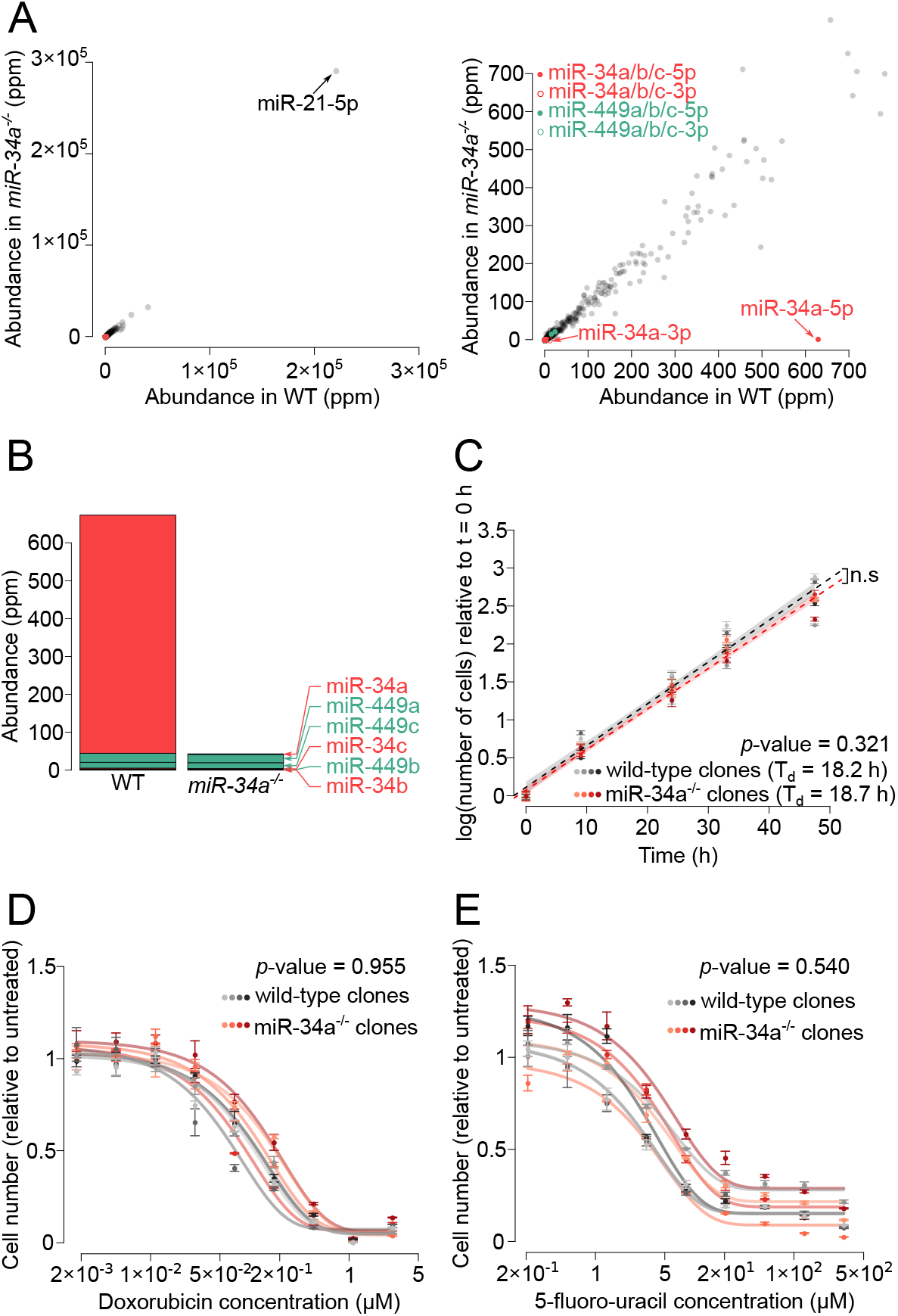
miR-34 is not a general repressor of cell proliferation. **A.** miRNA quantification by Small RNA-Seq in a representative wild-type HCT-116 clone *(x* axis) and a representative *miR-34a^−/−^* clone *(y* axis). Right panel: magnification of the left panel. **B.** Cumulated abundance of miR-34 family members in the two clones. miRNAs are sorted vertically according to their abundance in the wild-type clone. **C.** Four wild-type and four *miR-34a* mutant clones were grown in sub-confluent conditions. Means and standard errors of 4 biological replicates are represented by dots and error bars. Linear modeling of log-transformed cell counts relative to time was used to measure doubling time (T_d_), and to estimate the significance of the effect of genotype (p-value is given in the inset). Shaded areas represent the 95% confidence interval for theoretical future measurements. **D, E.** Cell number after 3 days of culture in presence of varying doses of **(D)** doxorubicin or **(E)** 5-fluoro-uracil (4 clones of each genotype were analyzed; 3 biological replicates for each drug concentration; mean +/− st. error is shown). Cell number was normalized to cell number count in untreated replicates. Normalized cell number was fitted to an asymptotic model for each clone (fitted models are represented by curves). In order to assess the significance of the effect of genotype, a naïve (non-informed by clone genotype) and a genotype-informed model were compared by an analysis of variance (p-value is indicated in the inset).

In agreement with published data, we did observe a strong reduction in cell proliferation when we transfected HCT-116 cells with large amounts (10 nM) synthetic miR-34a duplex (Figure 4**A**), but that effect was lost when transfecting 1 nM duplex (Figure 4**B**). Absolute miRNA quantification by RT-ddPCR shows that a 10 nM transfection over-expresses miR-34a by >8,000-fold in HCT-116 cells (and a 1 nM transfection over-expresses it by >490-fold), clearly demonstrating that such an experiment results in supra-physiological miRNA concentrations (Figure 4**C**). For comparison, we measured the increase in miR-34a expression in response to DNA damage: a 72 h treatment with doxorubicin at its IC50 concentration (7 × 10^-8^ M in HCT-116 cells; Supplementary Figure S7) over-expresses miR-34a by only 4.7-fold (Figure 4**D**).

**Figure 4:**
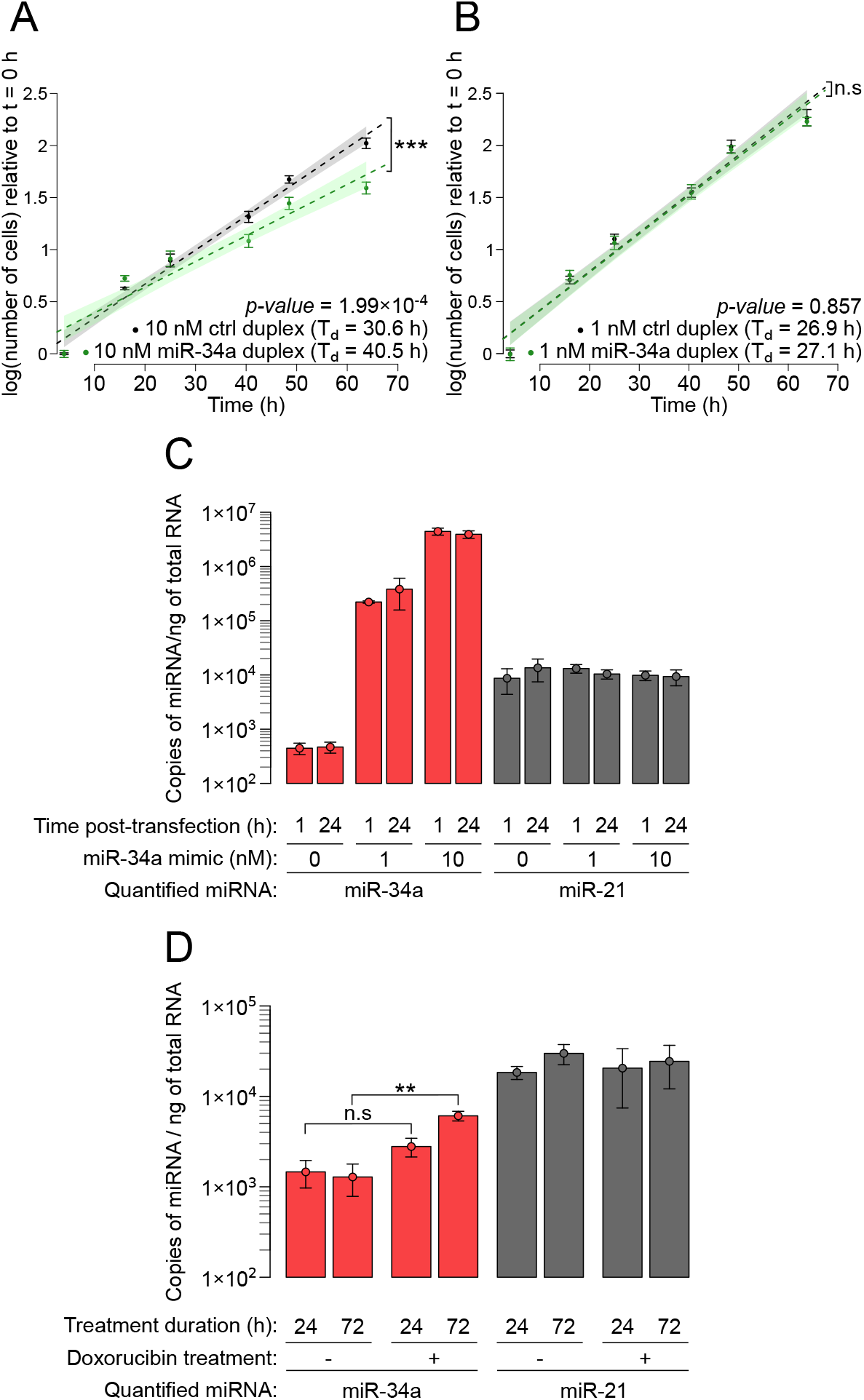
Supra-physiological transfection of miR-34a inhibits cell proliferation. Wild-type HCT-116 cells were transfected with 10 nM (panel **A**) or 1 nM (panel **B**) duplex (either a control siRNA duplex, or miR-34a/miR-34a* duplex) and grown in sub-confluent conditions. Means and standard errors of 6 biological replicates are represented by dots and error bars. Linear modeling of log-transformed cell counts relative to time was used to measure doubling time (T_d_), and to estimate the significance of the effect of duplex identity (p-values are given in the inset; asterisks denote *p-* value < 0.05, “n.s.” indicates larger p-values). Shaded areas represent the 95% confidence interval for theoretical future measurements. **C.** Cellular abundance of miR-34a (red bars) or a control miRNA (miR-21; gray bars) 1 or 24 h after transfection of HCT-116 cells with 0, 1 or 10 nM miR-34a/miR-34a* duplex. **D.** HCT-116 cells were treated for 24 or 72 h with 7 × 10^-8^ M doxorubicin, and their intracellular miR-34a and miR-21 were quantified by RT-ddPCR. Two-way ANOVA analysis shows that doxorubicin treatment has an effect on miR-34a levels (p=0.0013), and post-hoc pairwise t-tests find the effect significant only after 72 h exposure to the drug (p=0.0521 for 24 h exposure, p=0.00138 for 72 h exposure, indicated by “n.s.” and “**” respectively). A similar two-way ANOVA analysis does not detect a significant effect of doxorubicin treatment on miR-21 levels (p=0.768). **Panels C and D:** Means and standard deviations of 3 biological replicates are represented by dots and error bars, respectively.

It could be argued that low doses of transfected miR-34a could induce apoptosis, and our measurements of cell viability may have missed early apoptotic cells, therefore under-estimating the cytotoxicity of low dose miR-34a. This interpretation is ruled out by the measurement of Annexin V-labeled cells after transfection of various doses of miR-34a: physiological (picomolar range) doses do not appear to affect cell viability, and only the highest doses (>10 nM) lead to a visible decrease in cell viability (both through apoptosis and apoptosis-independent pathways): see Supplementary Figure S8.

## Discussion

Our results show that the *miR-34a* gene is rarely inactivated in cancers, whether by deletion, mutation or any other kind of process affecting miRNA expression. Its occasional loss in some cancers is most likely due to its genomic proximity with an actual tumor suppressor, and *miR-34a*-specific mutations are not enriched in any cancer type with data available in the largest cancer genomic dataset available. We also observed that the widely-assumed anti-proliferative role of miR-34a appears to be due to artifactual over-expression in cultured cells.

Of note, some authors have previously characterized the proliferative effect of miR-34 using genetic ablation rather than over-expression. In one study, mouse embryonic fibroblasts (MEFs) devoid of miR-34a/b/c appear to grow at the same rate than wild-type MEFs, except, transiently, for one early time-point [22]. Mutation of *miR-34a* alone also appeared not to affect MEF proliferation [32]. In another study, genetic inactivation of the *miR-34a* gene in HCT-116 is reported to accelerate cell proliferation, in stark contrast with our own findings [33]. Such discrepancy would deserve to be investigated, but unfortunately that published mutant cell line has been lost and it is no longer available from the authors (Dr. J. Lieberman, personal communication).

While the miR-34 family is believed to exert a tumor suppressive action in a diversity of cancers [21], we observed that it is hardly expressed in cultured cell lines, primary tissues and body fluids (Supplementary Figures S9-S11). It could be argued that a low level of miR-34 expression is expected in normal tissues, where p53 is mostly inactive. But p53 is clearly not the only regulator for miR-34, and the expression of miR-34 does not mirror p53 activity [22]. Current RNA detection technologies can be extremely sensitive, and they can detect miRNAs which are too poorly abundant to induce any clear change in target expression [34]. Hence we anticipate that in all the cell lines for which we analyzed miRNA abundance, and in most cells in the analyzed tissues, miR-34 family miRNAs are actually non-functional.

Yet we do not question the overall functionality of miR-34 miRNAs *in vivo.* Because that family is deeply conserved in evolution (shared between, *e.g.,* vertebrates and insects), it certainly plays important biological functions, perhaps only in a small number of cells, or at very specific developmental stages, where its abundance would be high enough. In mouse, the miR-34 family is particularly expressed in lungs and testes [22, 35]. Mutation of all 6 members of the miR-34 family causes severe ciliogenesis defects, leading to respiratory distress and impaired gametogenesis – translating into sterility and premature mortality [35]. Unsurprisingly then, the most obvious biological functions of that miRNA family seem to take place in the tissues where miR-34 miRNAs are highly expressed, in contrast with the widely-accepted notion of their broad anti-tumorigenic activity.

While the original definition for tumor suppressors had been formulated with coding genes in mind, we consider that there is no objective reason for adopting a different definition for tumor suppressor miRNAs. In this view, the most heavily studied candidate tumor suppressor miRNA, miR-34a, does not appear to be a tumor suppressor. It remains formally possible that miR-34a inactivation is frequent in specific cancer types, distinct from those we could analyze in Figures 1 and 2 and Supplementary Figure S2-3. In that case, miR-34a may be a tumor suppressor in these particular cancers, but rigorous investigation – while avoiding the pitfalls described above – of its impact on cell proliferation and tumorigenesis would be necessary to conclude so.

We confirmed that a large artificial over-expression (10 nM) of miR-34a indeed represses cell proliferation. It could be argued that this cytotoxic effect could provide the ground for an efficient anti-cancer treatment, no matter how un-natural it is. But the whole purpose of using natural tumor suppressors *(e.g.,* miRNAs) is that they are expected to be well tolerated, because they already exist endogenously. Administering large amounts of cytotoxic agents to patients may indeed kill cancer cells – but it will also likely trigger unwanted adverse effects. In this view, synthetic miR-34a behaves similarly to existing anti-cancer drugs, which are based on exogenous molecules. It is therefore not surprising to observe a variety of adverse secondary effects when the MRX34 miR-34a mimic is administered to patients [25, 24]. More inocuous miRNA-based treatments may be possible, but they would have to rely on rigorously established tumor-suppressive activity of the endogenous miRNA.

## Acknowledgments

We thank G. Canal, C. Theillet, P.D. Zamore and members of the Seitz lab for critical reading of the manuscript, and A. Pélisson and K. Mochizuki for useful discussions. This research was supported by Cancéropôle GSO “Émergence” grant, by Fondation ARC (grant “Projet” #PJA 20191209613 to the Seitz lab and PhD fellowship #ARCDOC42019120001002 to S.M.). We thank the MGX facility (Biocampus Montpellier, CNRS, INSERM, Univ. Montpellier, Montpellier, France) for sequencing the Small RNA-Seq libraries.

## Author contributions

S.M. and É.H. performed experiments; S.M. and H.S. performed computational analyses; S.M. and H.S. wrote the manuscript and prepared figures.

## References

[1] Green, A. R. (1988) Recessive mechanisms of malignancy. Br J Cancer, 58(2), 115–121.

[2] Weinberg, R. A. (1991) Tumor suppressor genes. Science, 254(5035), 1138–1146.

[3] Iwakawa, H. O. and Tomari, Y. (2015) The functions of MicroRNAs: mRNA decay and translational repression. Trends Cell Biol, 25(11), 651–665.

[4] Bartel, D. P. (2009) MicroRNAs: target recognition and regulatory functions. Cell, 136(2), 215–233.

[5] Friedman, R. C., Farh, K. K., Burge, C. B., and Bartel, D. P. (2009) Most mammalian mRNAs are conserved targets of microRNAs. Genome Res, 19(1), 92–105.

[6] Zhang, B., Pan, X., Cobb, G. P., and Anderson, T. A. (2007) microRNAs as oncogenes and tumor suppressors. Dev Biol, 302(1), 1–12.

[7] Wong, K. Y., Yu, L., and Chim, C. S. (2011) DNA methylation of tumor suppressor miRNA genes: a lesson from the miR-34 family. Epigenomics, 3(1), 83–92.

[8] Adams, B. D., Parsons, C., and Slack, F. J. (2016) The tumor-suppressive and potential therapeutic functions of miR-34a in epithelial carcinomas. Expert Opin Ther Targets, 20(6), 737–753.

[9] He, L., He, X., Lim, L. P., de Stanchina, E., Xuan, Z., Liang, Y., Xue, W., Zender, L., Magnus, J., Ridzon, D., Jackson, A. L., Linsley, P. S., Chen, C., Lowe, S. W., Cleary, M. A., and Hannon, G. J. (2007) A microRNA component of the p53 tumour suppressor network. Nature, 447(7148), 1130–1134.

[10] Bommer, G. T., Gerin, I., Feng, Y., Kaczorowski, A. J., Kuick, R., Love, R. E., Zhai, Y., Giordano, T. J., Qin, Z. S., Moore, B. B., MacDougald, O. A., Cho, K. R., and Fearon, E. R. (2007) p53-mediated activation of miRNA34 candidate tumor-suppressor genes. Curr Biol, 17(15), 1298–1307.

[11] Chang, T.-C., Wentzel, E. A., Kent, O. A., Ramachandran, K., Mullendore, M., Lee, K. H., Feldmann, G., Yamakuchi, M., Ferlito, M., Lowenstein, C. J., Arking, D. E., Beer, M. A., Maitra, A., and Mendell, J. T. (2007) Transactivation of miR-34a by p53 broadly influences gene expression and promotes apoptosis. Mol Cell, 26(5), 745–752.

[12] Corney, D. C., Flesken-Nikitin, A., Godwin, A. K., Wang, W., and Nikitin, A. Y. (2007) MicroRNA-34b and MicroRNA-34c are targets of p53 and cooperate in control of cell proliferation and adhesion-independent growth. Cancer Res, 67(18), 8433–8438.

[13] Tarasov, V., Jung, P., Verdoodt, B., Lodygin, D., Epanchintsev, A., Menssen, A., Meister, G., and Hermeking, H. (2007) Differential regulation of microRNAs by p53 revealed by massively parallel sequencing: miR-34a is a p53 target that induces apoptosis and G1-arrest. Cell Cycle, 6(13), 1586–1593.

[14] Tazawa, H., Tsuchiya, N., Izumiya, M., and Nakagama, H. (2007) Tumor-suppressive miR-34a induces senescence-like growth arrest through modulation of the E2F pathway in human colon cancer cells. Proc Natl Acad Sci USA, 104(39), 15472–15477.

[15] Raver-Shapira, N., Marciano, E., Meiri, E., Spector, Y., Rosenfeld, N., Moskovits, N., Bentwich, Z., and Oren, M. (2007) Transcriptional activation of miR-34a contributes to p53-mediated apoptosis. Mol Cell, 26(5), 731–743.

[16] Welch, C., Chen, Y., and Stallings, R. L. (2007) MicroRNA-34a functions as a potential tumor suppressor by inducing apoptosis in neuroblastoma cells. Oncogene, 26(34), 5017–5022.

[17] Lodygin, D., Tarasov, V., Epanchintsev, A., Berking, C., Knyazeva, T., Körner, H., Knyazev, P., Diebold, J., and Hermeking, H. (2008) Inactivation of miR-34a by aberrant CpG methylation in multiple types of cancer. Cell Cycle, 7(16), 2591–2600.

[18] Gallardo, E., Navarro, A., Viñolas, N., Marrades, R. M., Diaz, T., Gel, B., Quera, A., Bandres, E., Garcia-Foncillas, J., Ramirez, J., and Monzo, M. (2009) miR-34a as a prognostic marker of relapse in surgically resected non-small-cell lung cancer. Carcinogenesis, 30(11), 1903–1909.

[19] Wiggins, J. F., Ruffino, L., Kelnar, K., Omotola, M., Patrawala, L., Brown, D., and Bader, A. G. (2010) Development of a lung cancer therapeutic based on the tumor suppressor microRNA-34. Cancer Res, 70(14), 5923–5930.

[20] Corney, D. C., Hwang, C.-I., Matoso, A., Vogt, M., Flesken-Nikitin, A., Godwin, A. K., Kamat, A. A., Sood, A. K., Ellenson, L. H., Hermeking, H., and Nikitin, A. Y. (2010) Frequent downregulation of miR-34 family in human ovarian cancers. Clin Cancer Res, 16(4), 1119–1128.

[21] Slack, F. J. and Chinnaiyan, A. M. (2019) The Role of Non-coding RNAs in Oncology. Cell, 179(5), 1033–1055.

[22] Concepcion, C. P., Han, Y.-C., Mu, P., Bonetti, C., Yao, E., D’Andrea, A., Vidigal, J. A., Maughan, W. P., Ogrodowski, P., and Ventura, A. (2012) Intact p53-dependent responses in miR-34-deficient mice. PLoS Genet, 8(7), e1002797.

[23] Bader, A. G. (2012) miR-34 – a microRNA replacement therapy is headed to the clinic. Front Genet, 3, 120.

[24] Hong, D. S., Kang, Y.-K., Borad, M., Sachdev, J., Ejadi, S., Lim, H. Y., Brenner, A. J., Park, K., Lee, J.-L., Kim, T.-Y., Shin, S., Becerra, C. R., Falchook, G., Stoudemire, J., Martin, D., Kelnar, K., Peltier, H., Bonato, V., Bader, A. G., Smith, S., Kim, S., O’Neill, V., and Beg, M. S. (2020) Phase 1 study of MRX34, a liposomal miR-34a mimic, in patients with advanced solid tumours. Br J Cancer, 122(11), 1630–1637.

[25] Beg, M. S., Brenner, A. J., Sachdev, J., Borad, M., Kang, Y.-K., Stoudemire, J., Smith, S., Bader, A. G., Kim, S., and Hong, D. S. (2017) Phase I study of MRX34, a liposomal miR-34a mimic, administered twice weekly in patients with advanced solid tumors. Invest New Drugs, 35(2), 180–188.

[26] Concordet, J.-P. and Haeussler, M. (2018) CRISPOR: intuitive guide selection for CRISPR/Cas9 genome editing experiments and screens. Nucleic Acids Res, 46(W1), W242–W245.

[27] Ran, F. A., Hsu, P. D., Wright, J., Agarwala, V., Scott, D. A., and Zhang, F. (2013) Genome engineering using the CRISPR-Cas9 system. Nat Protoc, 8(11), 2281–2308.

[28] Kleinstiver, B. P., Pattanayak, V., Prew, M. S., Tsai, S. Q., Nguyen, N. T., Zheng, Z., and Joung, J. K. (2016) High-fidelity CRISPR-Cas9 nucleases with no detectable genome-wide off-target effects. Nature, 529(7587), 490–495.

[29] Zhang, Z., Hernandez, K., Savage, J., Li, S., Miller, D., Agrawal, S., Ortuno, F., Staudt, L. M., Heath, A., and Grossman, R. L. (2021) Uniform genomic data analysis in the NCI Genomic Data Commons. Nat Commun, 12(1), 1226.

[30] Bagchi, A., Papazoglu, C., Wu, Y., Capurso, D., Brodt, M., Francis, D., Bredel, M., Vogel, H., and Mills, A. A. (2007) *CHD5* is a tumor suppressor at human *1p36*. Cell, 128(3), 459–475.

[31] Mockly, S. and Seitz, H. (2019) Inconsistencies and Limitations of Current MicroRNA Target Identification Methods. Methods Mol Biol, 1970, 291–314.

[32] Choi, Y. J., Lin, C.-P., Ho, J. J., He, X., Okada, N., Bu, P., Zhong, Y., Kim, S. Y., Bennett, M. J., Chen, C., Ozturk, A., Hicks, G. G., Hannon, G. J., and He, L. (2011) miR-34 miRNAs provide a barrier for somatic cell reprogramming. Nat Cell Biol, 13(11), 1353–1360.

[33] Navarro, F. and Lieberman, J. (2015) miR-34 and p53: New Insights into a Complex Functional Relationship. PLoS One, 10(7), e0132767.

[34] Mullokandov, G., Baccarini, A., Ruzo, A., Jayaprakash, A. D., Tung, N., Israelow, B., Evans, M. J., Sachidanandam, R., and Brown, B. D. (2012) High-throughput assessment of microRNA activity and function using microRNA sensor and decoy libraries. Nat Methods, 9(8), 840–846.

[35] Song, R., Walentek, P., Sponer, N., Klimke, A., Lee, J. S., Dixon, G., Harland, R., Wan, Y., Lishko, P., Lize, M., Kessel, M., and He, L. (2014) miR-34/449 miRNAs are required for motile ciliogenesis by repressing cp110. Nature, 510(7503), 115–120.

