## Supplementary data for "A rationalized definition of general tumor suppressor microRNAs excludes miR-34a"

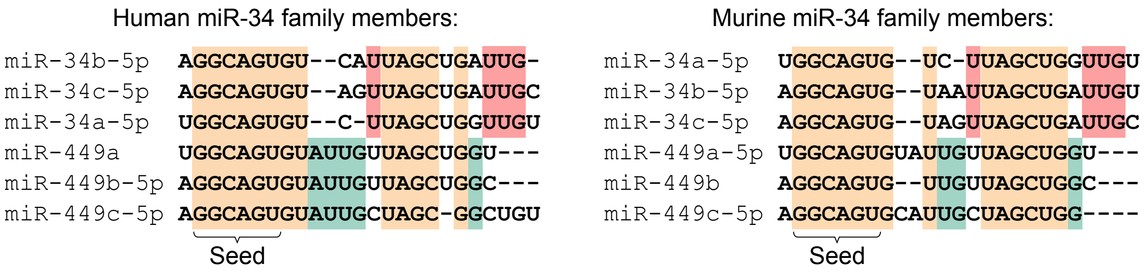


Supplementary Figure S 1: **Human and murine miR-34 family members.** Sequence alignment of the human (left) and murine (right) members of the miR-34 family. Nucleotides conserved between every family member are shown in orange. Nucleotides specific for the miR-34a/b/c subfamily are in red, those specific for the miR-449a/b/c subfamily are in green.

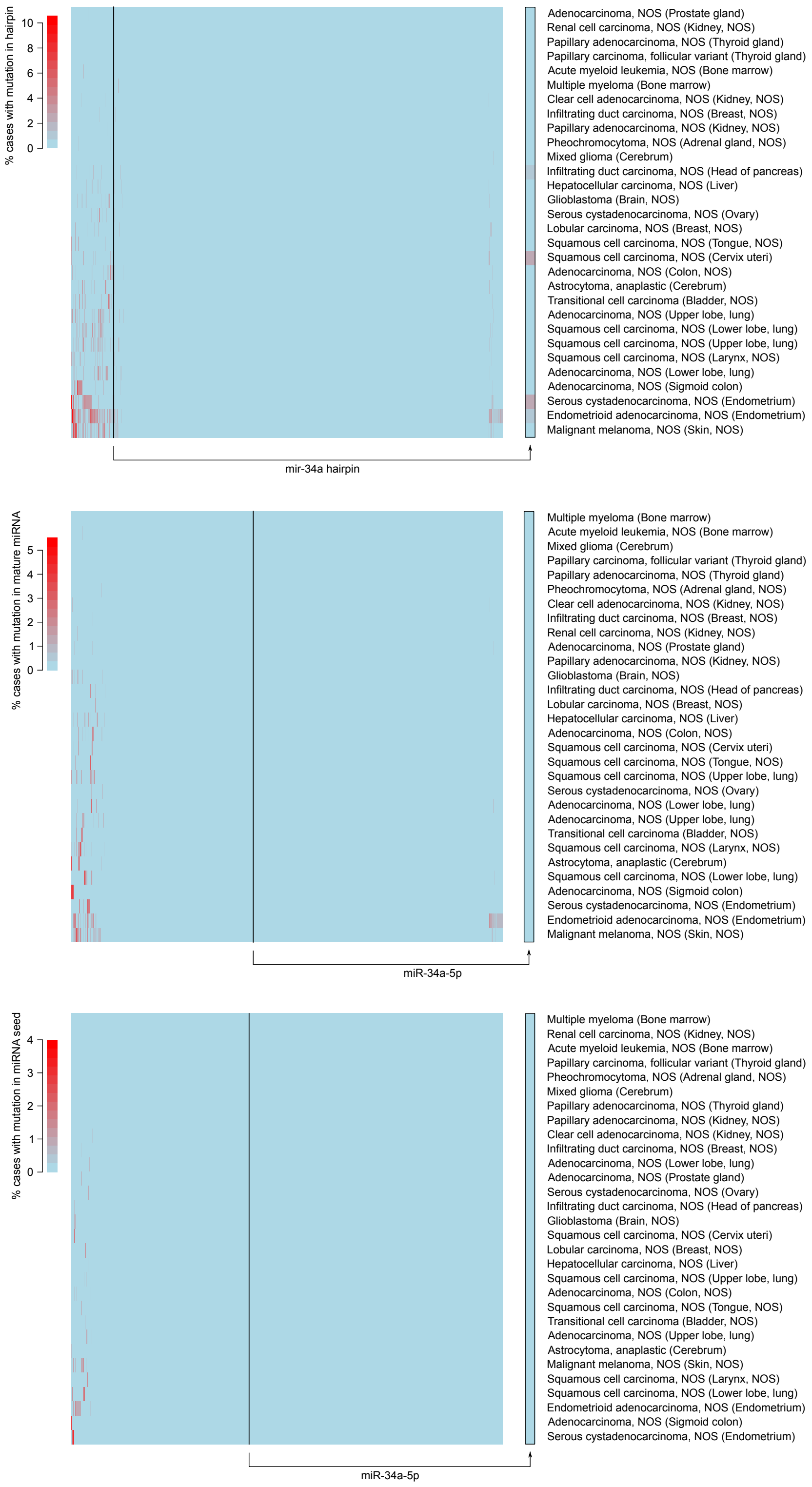


Supplementary Figure S2: ***mir-34a* is not generally mutated in cancers.** Only cancer types for which at least 100 cases were analyzed have been considered (n=30 cancer types; rows), and miRNA genes are shown in columns (n=1,750 hairpin loci in the top panel; 2,588 mature miRNA loci in the middle panel; 2,588 miRNA seed loci in the bottom panel). For each miRNA/cancer type pair, the heatmap shows the percentage of cases with recorded sequence variations either in the miRNA hairpin precursor (top panel), in the mature miRNA sequence (middle panel) or in the miRNA seed (bottom panel). miRNA loci were defined as in miRBase v.21. The column showing *mir-34a* data is magnified on the right margin (framed in black). “NOS”: not otherwise specified.

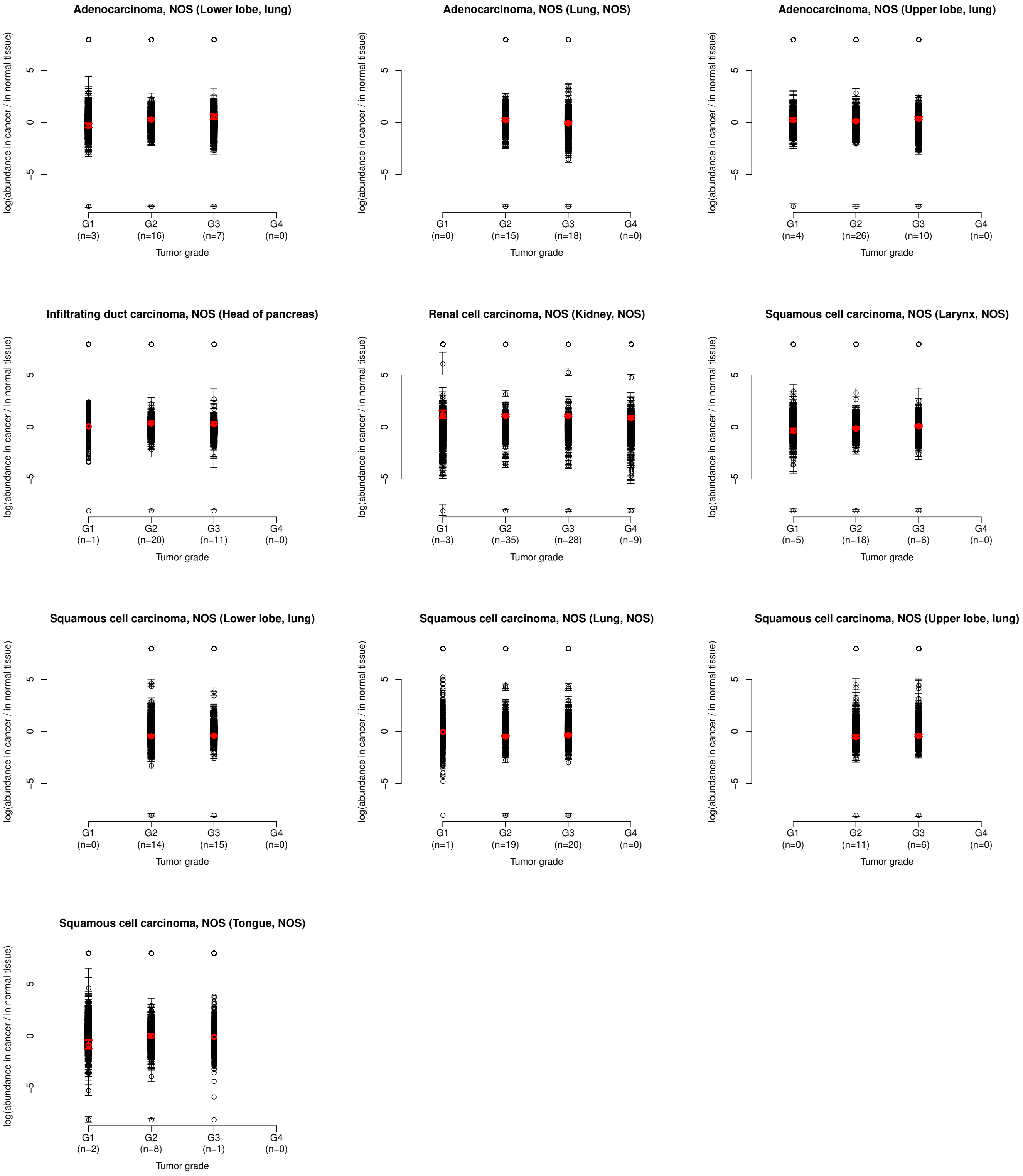


Supplementary Figure S3: **No evidence for *mir-34a* inactivation in the most aggressive tumors.** Cancer samples analyzed in Figure 1**A** were stratified by cancer grade (excluding cases whose grade was not determined, and selecting cancer types with at least 10 cases after this selection). miRNA abundance (normalized by the number of mapped miRNA reads) was compared between primary tumors and normal adjacent tissues. Each miRNA is represented by a dot (mean and standard error across independent cases are shown as a circle and error bars, respectively), with miR-34a being shown in red. log(fold-changes) larger than +8 or smaller than -8 were set to +8 or -8 respectively, for graphical clarity. "NOS": not otherwise specified. The number of analyzed cases is indicated under the $x$-axis for each tumor grade.

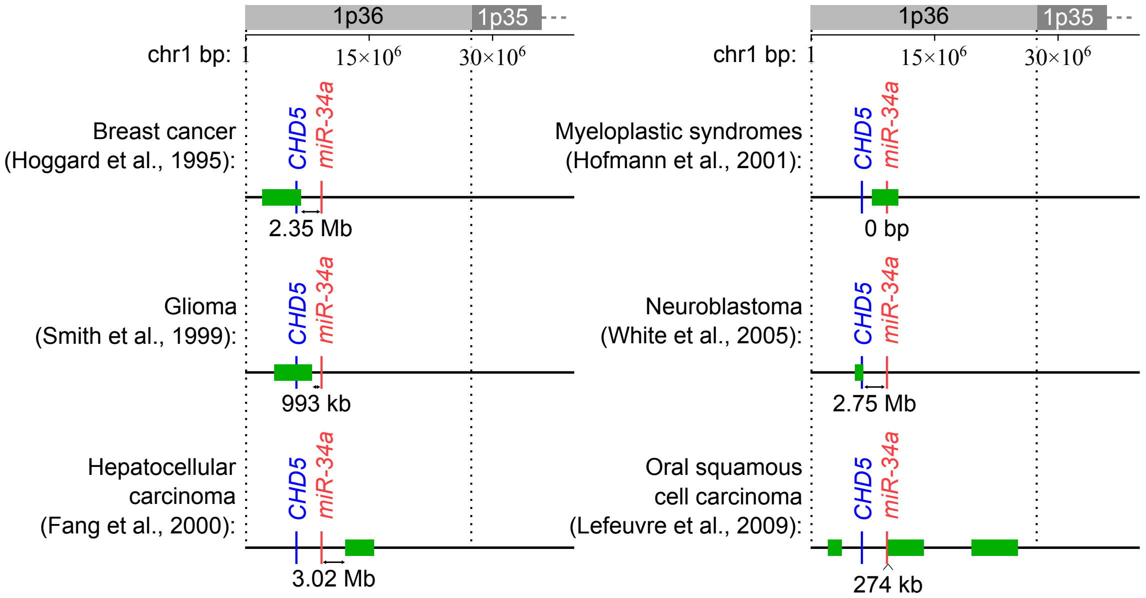


Supplementary Figure S4: **Smallest regions of consistent deletion identified in the 1p36 locus.** The 1p36 locus contains 885 annotated coding genes and 36 miRNA genes (Ensembl v.101); for clarity, only *CHD5* (in blue) and *miR-34a* (in red) are shown. Smallest identified region of consistent deletion are in green, and their distance to *miR-34* is indicated under the map. Genomic coordinates relate to the hg38 assembly.

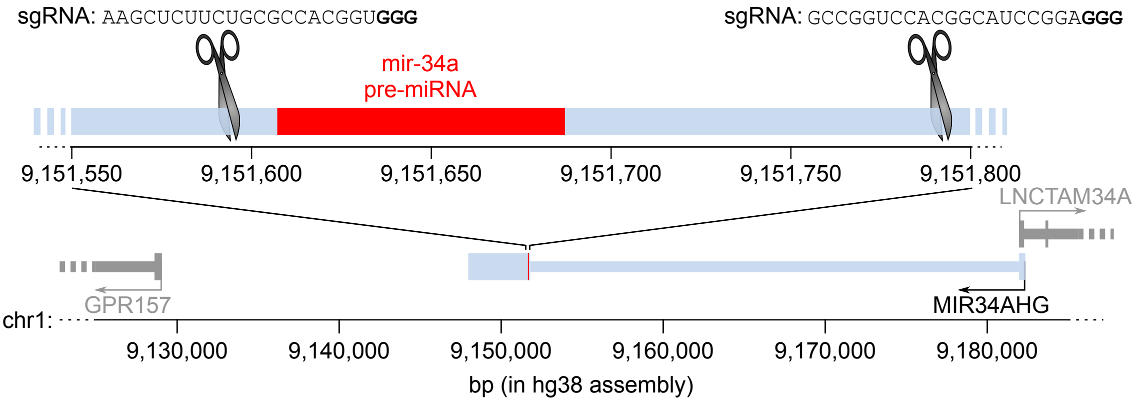


Supplementary Figure S5: **Strategy for CRISPR-mediated deletion of the *miR-34a* locus.** Thick lines: exons; thin lines: introns. Scissors represent Cas9-mediated cleavage sites, and the sequences of the cognate sgRNAs are written above each (PAM sequences in bold).

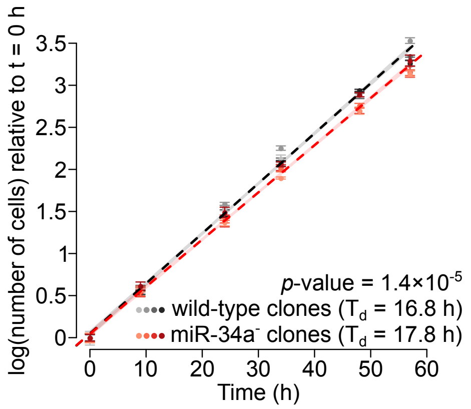


Supplementary Figure S6: **Proliferative effect of *miR-34a* in HAP1 cells.** Four wild-type and four *miR-34a* mutant clones were grown in sub-confluent conditions. Means and standard errors of 4 biological replicates are represented by dots and error bars. Linear modeling of log-transformed cell counts relative to time was used to measure doubling time ($T_d$), and to estimate the significance of the effect of genotype ($p$-value is given in the inset). Shaded areas represent the 95% confidence interval for theoretical future measurements.

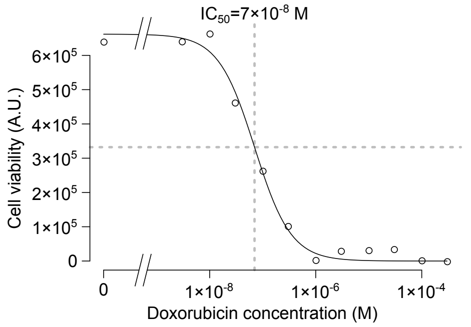


Supplementary Figure S7: **Determination of the half-maximal inhibitory concentration ($IC_{50}$) of doxorubicin in HCT-116 cells.** Wild-type HCT-116 cells were grown in various concentrations of doxorubicin (shown on the $x$-axis), and cell viability was measured after 72 h using by ATP quantification (results shown on the $y$-axis).

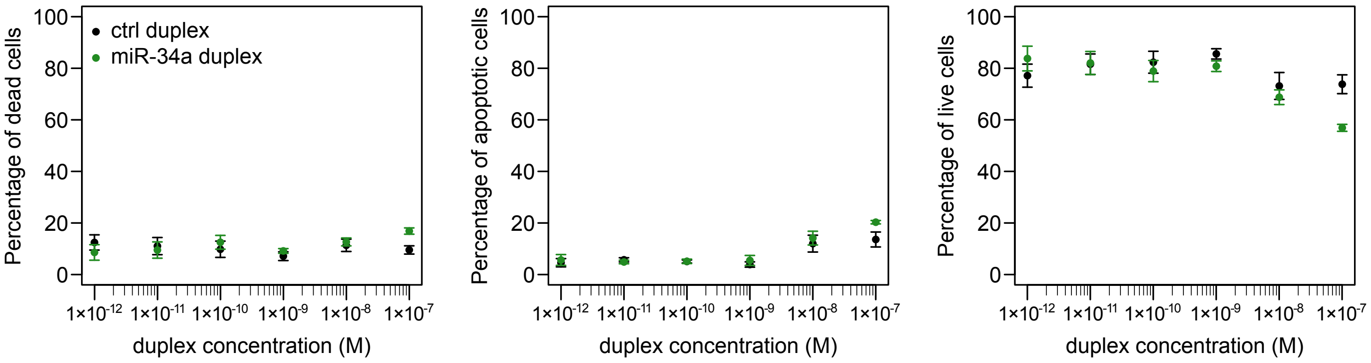


| **Two-way ANOVA *p*-value** | dead | apoptotic | alive |
|---|---|---|---|
| duplex concentration | 0.465 | $1.02 \times 10^{-7}$ | $5.84 \times 10^{-4}$ |
| duplex identity | 0.365 | 0.104 | 0.129 |

Supplementary Figure S8: **No effect of sub-nanomolar doses of miR-34a on HCT-116 cell death or apoptosis.** Cell viability and apoptosis were assessed 72 h after transfection of various concentrations of control (black) or miR-34a (green) duplex, using FACS analysis after FITC Annexin V- and PI-staining. Three biological replicates were performed, means are represented by dots and standard errors are represented by error bars. Left, middle and right panels represent the percentage of dead (FITC Annexin V-negative and PI-positive), apoptotic (FITC Annexin V-positive and PI-negative) and live (FITC Annexin V-negative and PI-negative) cells respectively. The table in the bottom panel indicates *p*-values for a two-way ANOVA test without interaction interrogating the effect of duplex concentration and duplex identity on the percentage of each category (dead, apoptotic, or alive).

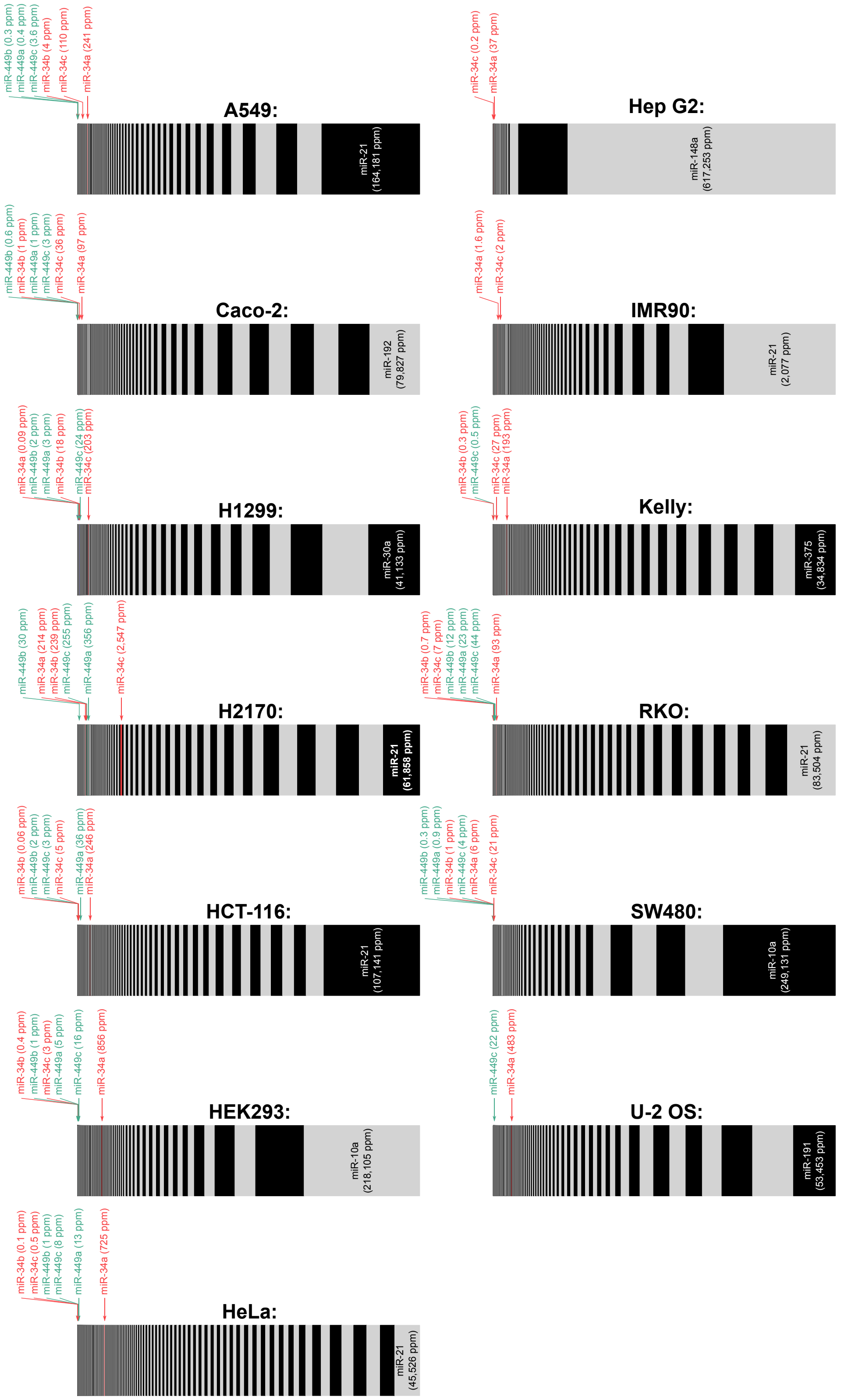


Supplementary Figure S9: **miRNA abundance in human cell lines.** miRNAs are ranked by increasing abundance from left to right, and the width of each rectangle is proportional to miRNA abundance. Members of the miR-34 family, and their expression level, are shown in red and green (red for the miR-34a/b/c subfamily, green for the miR-449a/b/c subfamily). miRNA abundance is normalized to the total number of genome-matching reads, and expressed as *parts per million* (ppm). Small RNA-Seq datasets used in this figure are listed in Supplementary Table 2.

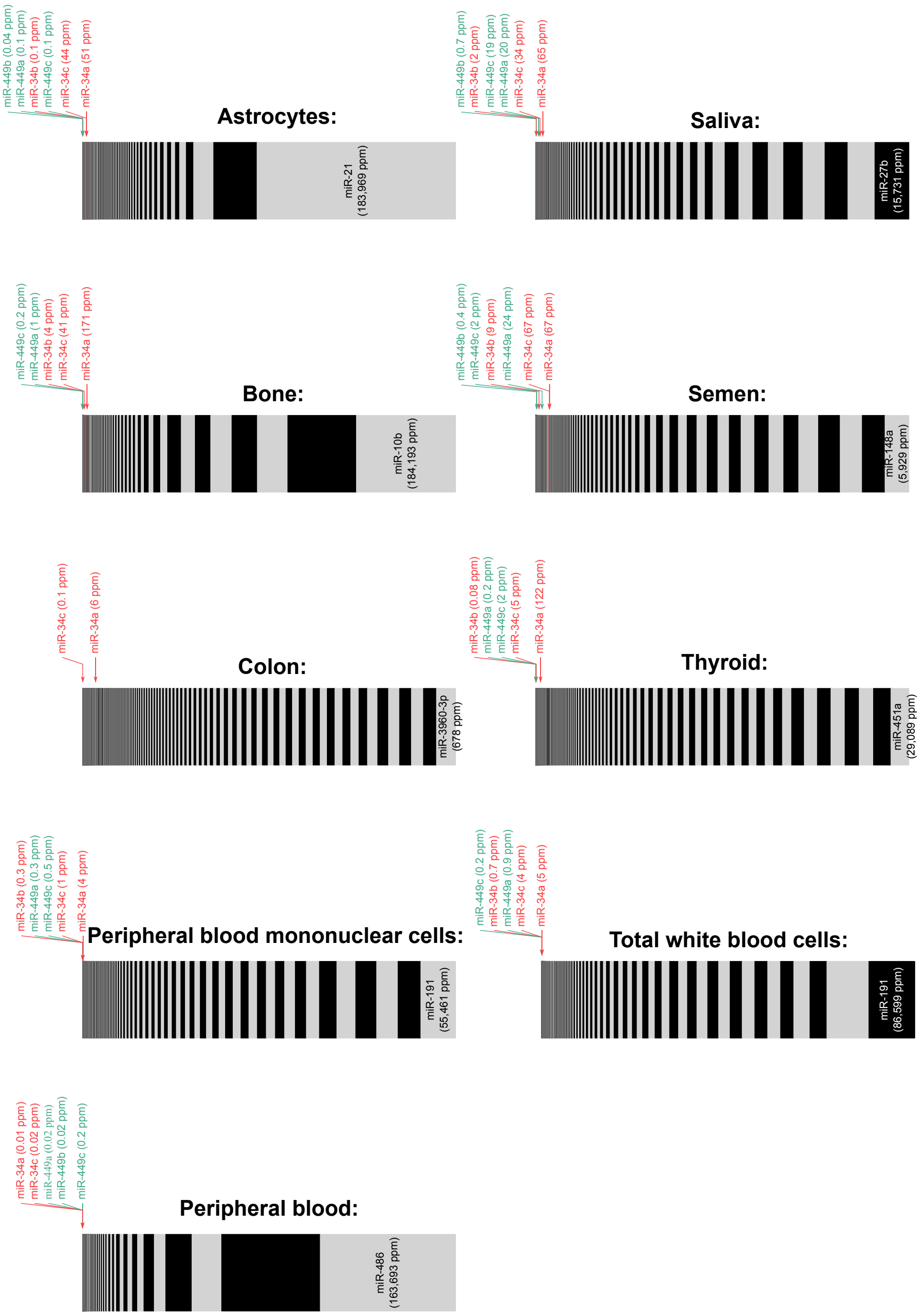


Supplementary Figure S10: **miRNA abundance in human tissues and body fluids.** miRNAs are ranked by increasing abundance from left to right, and the width of each rectangle is proportional to miRNA abundance. Members of the miR-34 family, and their expression level, are shown in red and green (red for the miR-34a/b/c subfamily, green for the miR-449a/b/c subfamily). miRNA abundance is normalized to the total number of genome-matching reads, and expressed as *parts per million* (ppm). Small RNA-Seq datasets used in this figure are listed in Supplementary Table 3.

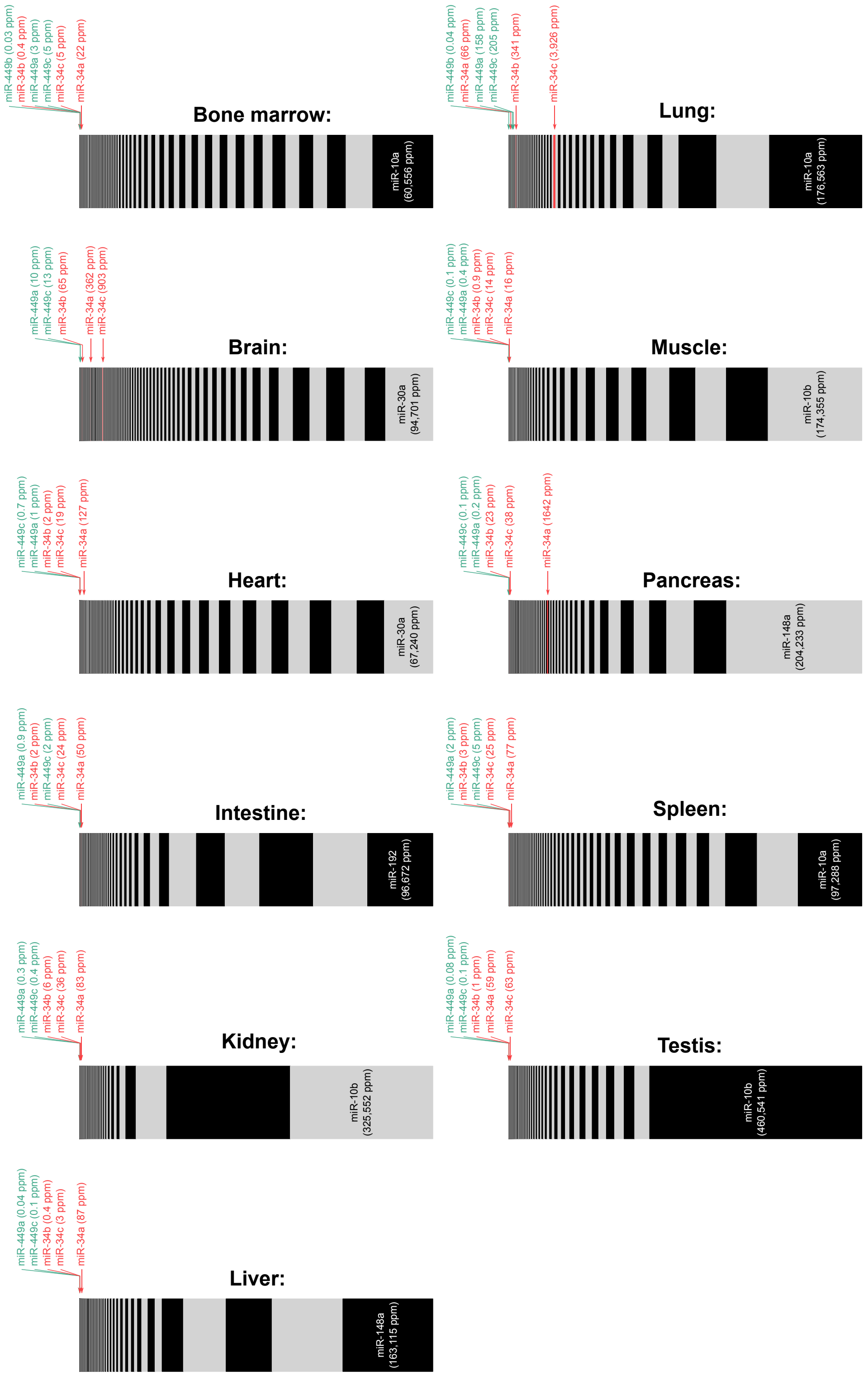


Supplementary Figure S11: **miRNA abundance in murine tissues.** miRNAs are ranked by increasing abundance from left to right, and the width of each rectangle is proportional to miRNA abundance. Members of the miR-34 family, and their expression level, are shown in red and green (red for the miR-34a/b/c subfamily, green for the miR-449a/b/c subfamily). miRNA abundance is normalized to the total number of genome-matching reads, and expressed as *parts per million* (ppm). Small RNA-Seq datasets used in this figure are listed in Supplementary Table 4.

| miRNA gene | Confidence |
|---|---|
| mir-4767 | lower |
| mir-4770 | lower |
| mir-651 | high |
| mir-548ax | lower |
| mir-3915 | lower |
| mir-548f-5 | lower |
| mir-6134 | lower |
| mir-4666b | lower |
| mir-23c | lower |
| mir-548am | lower |
| mir-4768 | lower |
| mir-548aj-2 | lower |
| mir-1587 | lower |
| mir-3937 | lower |
| mir-4769 | lower |
| mir-222 | high |
| mir-221 | high |
| mir-98 | high |
| let-7f-2 | high |
| mir-6857 | lower |
| mir-6895 | lower |
| mir-6894 | lower |
| mir-8088 | lower |
| mir-502 | lower |
| mir-660 | high |
| mir-500a | high |
| mir-532 | high |
| mir-501 | lower |
| mir-500b | lower |
| mir-362 | high |
| mir-188 | high |
| mir-1468 | lower |
| mir-223 | high |
| mir-361 | high |
| mir-548i-4 | lower |

| miRNA gene | Confidence |
|---|---|
| mir-325 | lower |
| mir-545 | high |
| mir-374a | high |
| mir-421 | lower |
| mir-374b | high |
| mir-1184-3 | lower |
| mir-1184-2 | lower |
| mir-1184-1 | lower |
| mir-664b | high |
| mir-6858 | lower |
| mir-3202-1 | lower |
| mir-718 | lower |
| mir-6087 | lower |
| mir-548m | lower |
| mir-3672 | lower |
| mir-766 | high |
| mir-1277 | lower |
| mir-548an | lower |
| mir-3978 | lower |
| mir-652 | high |
| mir-4329 | lower |
| mir-1912 | lower |
| mir-764 | lower |
| mir-448 | lower |
| mir-1298 | lower |
| mir-1911 | lower |
| mir-320d-2 | lower |
| mir-504 | high |
| mir-934 | lower |
| mir-505 | high |
| mir-92a-2 | high |
| mir-106a | high |
| mir-363 | high |
| mir-19b-2 | high |
| mir-20b | high |

| miRNA gene | Confidence |
|---|---|
| mir-542 | high |
| mir-450a-2 | high |
| mir-424 | high |
| mir-450b | high |
| mir-450a-1 | high |
| mir-503 | high |
| mir-4330 | lower |
| mir-452 | lower |
| mir-224 | lower |
| mir-767 | high |
| mir-105-1 | high |
| mir-105-2 | high |
| mir-891a | lower |
| mir-892c | lower |
| mir-891b | lower |
| mir-892a | lower |
| mir-888 | lower |
| mir-890 | lower |
| mir-892b | lower |
| mir-2114 | lower |
| mir-514a-1 | high |
| mir-514a-3 | high |
| mir-514a-2 | high |
| mir-510 | high |
| mir-509-3 | high |
| mir-509-1 | high |
| mir-513b | lower |
| mir-513c | high |
| mir-513a-1 | lower |
| mir-513a-2 | lower |
| mir-514b | high |
| mir-508 | high |
| mir-509-2 | high |
| mir-507 | lower |
| mir-506 | high |

Supplementary Table S1: **Frequently deleted miRNA genes in cancer.** Identity of the 105 miRNA genes frequently deleted in a variety of cancers (red area at the top left corner of Figure 1**B**). For each miRNA gene, its confidence level (as defined by miRBase v.21; [7]) is indicated.

| Cell line: | SRA accession number(s): |
|---|---|
| A549 | SRR6713501,SRR5689166, DRR036695 and SRR1304309 |
| Caco-2 | ERR3415707 |
| H1299 | DRR036697 |
| H2170 | SRR3341761 and SRR3341762 |
| HCT-116 | SRR954987, SRR954996, SRR4235725, SRR4235726, ERR3173398, ERR3173397 and ERR3173396 |
| HEK293 | SRR1240816 and SRR1240817 |
| HeLa | SRR8311268 and SRR8311269 |
| Hep G2 | SRR12054851 |
| IMR90 | SRR020286 |
| Kelly | SRR3533075, SRR3533074 and SRR3533073 |
| RKO | ERR3415712 |
| SW480 | SRR3923807 and SRR3923808 |
| U-2 OS | SRR10225092 |

Supplementary Table S2: **Small RNA-Seq datasets used in Supplementary Figure S9.** Several of these cell lines had been used in previous studies to evaluate the proliferative effect of miR-34 by over-expression experiments: A549 [8], H1299 [9], HCT-116 [8, 10, 11], Kelly [12], IMR90 [8], RKO [9], SW480 [13] and U-2 OS [9].

| Tissue or body fluid: | SRA accession numbers: |
|---|---|
| Astrocytes | SRR2915342, SRR2915343 and SRR2915344 |
| Bone | SRR6324194 |
| Colon | SRR6895202, SRR6895203, SRR6895204 and SRR6895205 |
| Peripheral blood mononuclear cells | SRR7412273–SRR7412275, SRR7412278, SRR7412280–SRR7412285, SRR7412296–SRR7412300, SRR7412302–SRR7412311, SRR7412313–SRR7412315 and SRR7412326–SRR7412334 |
| Peripheral blood | SRR9844335–SRR9844346, SRR9844348, SRR9844349, SRR9844351–SRR9844360, SRR9844362, SRR9844364, SRR9844366–SRR9844373, SRR9844375 and SRR9844377–SRR9844386 |
| Saliva | SRR3144036–SRR3144041, SRR3144044–SRR3144053, SRR3144055, SRR3144057 and SRR3144060 |
| Semen | SRR11912557–SRR11912563 and SRR11912574 |
| Thyroid | SRR8393464–SRR8393466 |
| Total white blood cells | SRR7012343 |

Supplementary Table S3: **Small RNA-Seq datasets used in Supplementary Figure S10.**

| **Tissue:** | **SRA accession numbers:** |
|---|---|
| Bone marrow | SRR10695972–SRR10695983 and SRR7807316–SRR7807327 |
| Brain | SRR10695984–SRR10695997 and SRR7807328–SRR7807341 |
| Heart | SRR10695998–SRR10696010 and SRR7807342–SRR7807354 |
| Intestine | SRR10696011–SRR10696022 and SRR7807355–SRR7807366 |
| Kidney | SRR10695896–SRR10695903, SRR10696023–SRR10696028<br>SRR7807237–SRR7807244 and SRR7807367–SRR7807372 |
| Liver | SRR10695904–SRR10695917 and SRR7807245–SRR7807258 |
| Lung | SRR10695918–SRR10695930 and SRR7807259–SRR7807271 |
| Muscle | SRR10695931–SRR10695944 and SRR7807272–SRR7807285 |
| Pancreas | SRR10695945–SRR10695957 and SRR7807286–SRR7807298 |
| Spleen | SRR10695958–SRR10695971 and SRR7807299–SRR7807312 |
| Testis | SRR10662083–SRR10662085 and SRR7807313–SRR7807315 |

Supplementary Table S4: **Small RNA-Seq datasets used in Supplementary Figure S11.**

### References


[1] Hoggard, N., Brintnell, B., Howell, A., Weissenbach, J., and Varley, J. (1995) Allelic imbalance on chromosome 1 in human breast cancer. II. Microsatellite repeat analysis. *Genes Chromosomes Cancer,* **12**(1), 24–31.

[2] Smith, J. S., Alderete, B., Minn, Y., Borell, T. J., Perry, A., Mohapatra, G., Hosek, S. M., Kimmel, D., O'Fallon, J., Yates, A., Feuerstein, B. G., Burger, P. C., Scheithauer, B. W., and Jenkins, R. B. (1999) Localization of common deletion regions on 1p and 19q in human gliomas and their association with histological subtype. *Oncogene,* **18**(28), 4144–4152.

[3] Fang, W., Piao, Z., Simon, D., Sheu, J. C., and Huang, S. (2000) Mapping of a minimal deleted region in human hepatocellular carcinoma to 1p36.13-p36.23 and mutational analysis of the *RIZ* (*PRDM2*) gene localized to the region. *Genes Chromosomes Cancer,* **28**(3), 269–275.

[4] Hofmann, W. K., Takeuchi, S., Xie, D., Miller, C. W., Hoelzer, D., and Koeffler, H. P. (2001) Frequent loss of heterozygosity in the region of D1S450 at 1p36.2 in myelodysplastic syndromes. *Leuk Res,* **25**(10), 855–858.

[5] White, P. S., Thompson, P. M., Gotoh, T., Okawa, E. R., Igarashi, J., Kok, M., Winter, C., Gregory, S. G., Hogarty, M. D., Maris, J. M., and Brodeur, G. M. (2005) Definition and characterization of a region of 1p36.3 consistently deleted in neuroblastoma. *Oncogene,* **24**(16), 2684–2694.

[6] Lefeuvre, M., Gunduz, M., Nagatsuka, H., Gunduz, E., Ali, M. A. S., Beder, L., Fukushima, K., Yamanaka, N., Shimizu, K., and Nagai, N. (2009) Fine deletion analysis of 1p36 chromosomal region in oral squamous cell carcinomas. *J Oral Pathol Med,* **38**(1), 94–98.

[7] Kozomara, A. and Griffiths-Jones, S. (2014) miRBase: annotating high confidence microRNAs using deep sequencing data. *Nucleic Acids Res,* **42**(Database issue), D68–D73.

[8] He, L., He, X., Lim, L. P., de Stanchina, E., Xuan, Z., Liang, Y., Xue, W., Zender, L., Magnus, J., Ridzon, D., Jackson, A. L., Linsley, P. S., Chen, C., Lowe, S. W., Cleary, M. A., and Hannon, G. J. (2007) A microRNA component of the p53 tumour suppressor network. *Nature,* **447**(7148), 1130–1134.

[9] Tarasov, V., Jung, P., Verdoodt, B., Lodygin, D., Epanchintsev, A., Menssen, A., Meister, G., and Hermeking, H. (2007) Differential regulation of microRNAs by p53 revealed by massively parallel sequencing: miR-34a is a p53 target that induces apoptosis and G1-arrest. *Cell Cycle,* **6**(13), 1586–1593.

[10] Chang, T.-C., Wentzel, E. A., Kent, O. A., Ramachandran, K., Mullendore, M., Lee, K. H., Feldmann, G., Yamakuchi, M., Ferlito, M., Lowenstein, C. J., Arking, D. E., Beer, M. A., Maitra, A., and Mendell, J. T. (2007) Transactivation of miR-34a by p53 broadly influences gene expression and promotes apoptosis. *Mol Cell,* **26**(5), 745–752.

[11] Tazawa, H., Tsuchiya, N., Izumiya, M., and Nakagama, H. (2007) Tumor-suppressive miR-34a induces senescence-like growth arrest through modulation of the E2F pathway in human colon cancer cells. *Proc Natl Acad Sci USA,* **104**(39), 15472–15477.

[12] Welch, C., Chen, Y., and Stallings, R. L. (2007) MicroRNA-34a functions as a potential tumor suppressor by inducing apoptosis in neuroblastoma cells. *Oncogene,* **26**(34), 5017–5022.

[13] Bommer, G. T., Gerin, I., Feng, Y., Kaczorowski, A. J., Kuick, R., Love, R. E., Zhai, Y., Giordano, T. J., Qin, Z. S., Moore, B. B., MacDougald, O. A., Cho, K. R., and Fearon, E. R. (2007) p53-mediated activation of miRNA34 candidate tumor-suppressor genes. *Curr Biol,* **17**(15), 1298–1307.
